# Systems-ecology designed bacterial consortium protects from severe *Clostridioides difficile* infection

**DOI:** 10.1101/2023.08.08.552483

**Authors:** Matthew L Jenior, Jhansi L Leslie, Glynis L Kolling, Laurie Archbald-Pannone, Deborah A Powers, William A Petri, Jason A Papin

**Affiliations:** Department of Biomedical Engineering, University of Virginia, Charlottesville, VA, USA; Department of Internal Medicine, Division of Infectious Diseases & International Health, University of Virginia, Charlottesville, VA, USA; Department of Biochemistry & Molecular Genetics, University of Virginia, Charlottesville, VA, USA; Department of Microbiology, Immunology and Cancer Biology, University of Virginia Health System, Charlottesville, Virginia, USA; Department of Pathology, University of Virginia Health System, Charlottesville, Virginia, USA; Department of Internal Medicine, University of Virginia Health System, Charlottesville, Virginia, USA; Department of Internal Medicine, Division of General, Geriatric, Palliative, and Hospital Medicine, University of Virginia School of Medicine, Charlottesville, VA, USA

## Abstract

Fecal Microbiota Transplant (FMT) is an emerging therapy that has had remarkable success in treatment and prevention of recurrent *Clostridioides difficile* infection (rCDI). FMT has recently been associated with adverse outcomes such as inadvertent transfer of antimicrobial resistance, necessitating development of more targeted bacteriotherapies. To address this challenge, we developed a novel systems biology pipeline to identify candidate probiotic strains that would be predicted to interrupt *C. difficile* pathogenesis. Utilizing metagenomic characterization of human FMT donor samples, we identified those metabolic pathways most associated with successful FMTs and reconstructed the metabolism of encoding species to simulate interactions with *C. difficile*. This analysis resulted in predictions of high levels of cross-feeding for amino acids in species most associated with FMT success. Guided by these *in silico* models, we assembled consortia of bacteria with increased amino acid cross-feeding which were then validated *in vitro*. We subsequently tested the consortia in a murine model of CDI, demonstrating total protection from severe CDI through decreased toxin levels, recovered gut microbiota, and increased intestinal eosinophils. These results support the novel framework that amino acid cross-feeding is likely a critical mechanism in the initial resolution of CDI by FMT. Importantly, we conclude that our predictive platform based on predicted and testable metabolic interactions between the microbiota and *C. difficile* led to a rationally designed biotherapeutic framework that may be extended to other enteric infections.

## INTRODUCTION

Fecal Microbial Transplant (FMT) refers to the transfer of healthy donor fecal material to the intestine of a recipient to alter their gut microbiome, conferring a health benefit^7–91^. The microbial ecosystem of the intestines is involved in several aspects of our physiology including educating the immune system, directly protecting from infection, regulating epithelial homeostasis, and augmenting host metabolism^1^. FMT has been effective to varying degrees across numerous inflammatory, neurological, and metabolic disorders with connections to the gut microbiota^2^. However, manipulation of the gut microbiome through FMT has proven most effective in treating recurrent *Clostridioides difficile* infection (rCDI). Unfortunately, in addition to positive effects, there have also been adverse health events including transfer of pathogens and inflammatory agents^3^.

CDI is the most frequent hospital-acquired infection in the United States and Europe^4–6^. Antibiotics are a major risk factor for contracting CDI, and many groups have shown that antibiotic treatment disrupts the gastrointestinal microbiota, altering host immunity, enabling colonization by *C. difficile*, and also increase the likelihood of recurrence^7–9^. FMT has been extremely effective for quickly resolving >90% of rCDI cases after a single infusion underscoring the importance of the gut microbial community in this infection ^10^. Metabolomic analyses of samples taken from FMT recipients revealed significant shifts in concentration of *C. difficile* growth substrates as well as germination signals including bile acids^11^. Additionally, it has also been shown that access to simple carbohydrates and fermentable amino acids downregulates *C. difficile* virulence expression^12–14^. Together, these findings support the hypothesis that bacterial co-metabolism is important to the mechanism of successful FMT and that microbial community members may possess cross-feeding metabolic interactions with *C. difficile* that lead to symptom resolution. This hypothesis is largely contrary to efforts where the prevailing theory has been that competition for nutrients is the primary mechanism behind microbe-mediated CDI resolution^15^.

Systems biology approaches have been effective for predicting specific mechanisms in complex biological phenotypes. Culture-based approaches have identified resident bacterial members of the murine microbiota that directly impact survival during CDI in gnotobiotic mice^16^. One set of computational tools for understanding metabolism within systems biology are genome-scale metabolic network reconstructions (GENREs). GENREs have recently been utilized with metagenomic sequencing from Crohn’s disease patients to uncover metabolic interactions between microbes that may influence disease^17,18^. Recently, we generated new GENREs for *C. difficile* to identify previously unappreciated metabolic links to virulence^19^. Since rCDI is caused by a single organism, and susceptibility is strongly associated with the gut microbiome, rCDI presents a unique opportunity for development of defined consortia based on metabolic pathways as replacement for FMT. We created a combined pipeline of *in silico*, *in vitro*, and *in vivo* methods to design and test a targeted consortia of bacterial species that metabolically interact with *C. difficile* during infection. By performing deep metagenomic sequencing on human FMT donor samples associated with both successful and failed resolution of rCDI, we found amino acid metabolism pathways were more predictive than species abundances in successful FMT. Isolating those metagenome-assembled genomes (MAGs) enriched for this functionality, we identified 26 MAGs for candidate probiotics which were used to generate GENREs to simulate both competitive and cooperative metabolic interactions with *C. difficile*. This analysis revealed that potential probiotic species with the greatest impact on *C. difficile* GENRE growth most frequently shared or competed for amino acids, which we subsequently validated *in vitro*. These results led to the creation of two four-member bacterial consortia that favored either high levels of metabolic cooperation or competition with *C. difficile*. We tested the effect of each consortium in an animal model of CDI. Administration of the cooperative consortium led to survival of all animals, while animals in both the competitive consortium and control groups succumbed to the infection. We found that rather than diminishing pathogen burden, treatment with the cooperative consortia significantly decreased *C. difficile* toxin levels, increased community diversity, and increased eosinophil abundance. These findings represent a paradigm shift in the thinking behind mechanisms of FMT efficacy; namely, that metabolic cooperation with *C. difficile* to downregulate virulence during healthy community recovery may outweigh the importance of outcompeting the pathogen for nutrients. This is the first instance that a computationally selected and designed live biotherapeutic, leveraging sequencing data from patient samples, has been created and shown efficacy in a pre-clinical animal model of disease.

## RESULTS

### Metagenomics from successful FMT supports the importance of metabolic interactions with *C. difficile*

To profile the bacterial taxa and metabolic pathways that we predicted would be overrepresented in successful FMT, we deeply sequenced metagenomes from a unique FMT donor cohort from a recent clinical study at the University of Virginia^20^. A total of 119 patients with confirmed rCDI were treated with FMT and followed to determine efficacy of the therapy. A failed FMT was defined as recurrence of diarrhea with a positive stool ELISA for *C. difficile* toxin or positive *tcdB* PCR within three months of FMT. Among the donor samples, we selected three individuals based on dissimilar rates of successful rCDI resolution across recipients (Fig. 1A). Donor A had a donation pool of 33 samples which were associated with a 90.9% success rate across patient recipients (30/33), while donors B and C had success rates of 53.3% (16/30) and 57.1% (8/14) respectively. From each donor, four samples were chosen which were reflective of that donor’s overall success rate, were separate donations, and went to a distinct recipient. From the 12 samples, shotgun metagenomic sequencing resulted in 158 million curated reads per sample (Table S1). This study design allowed for partial control for within individual variation among the donors as well as recipient-specific effects that may have impacted FMT efficacy. By focusing downstream analyses on donor A versus donors B & C combined, we sought to identify those microbial features that most associated with consistent successful FMT.

**Figure 1:**
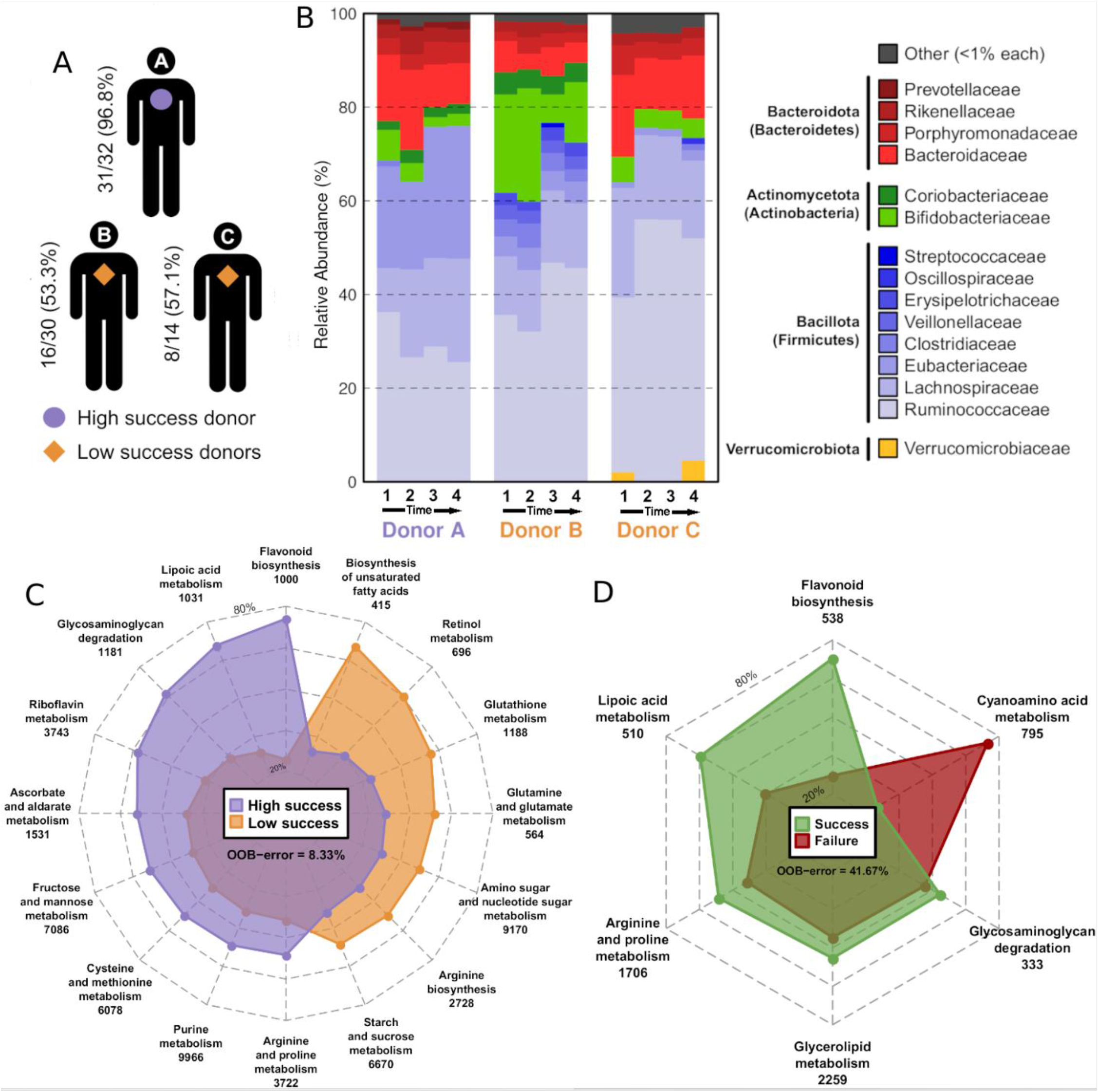
Shotgun metagenomic analysis from FMT donors in local patient cohort. **(A)** Human FMT sample schematic, following previous FMT study at UVA. Three individual donors with >14 donations each were selected for metagenomic profiling based on rates of successful rCDI resolution. Donor A had the highest rate of success with 30 of 33 donations resulting in resolved rCDI (90.9%). The other two donors had rates of Donor B 16 of 30 donations (53.3%) and Donor C 8 of 14 samples (57.1%), respectively. **(B)** Predicted Family-level classification using MetaPhlAn2. Donors, time, and associated outcomes labeled on lower axis according to panel A. **(C)** Superorganism-level analysis of metabolic capacity present in the microbiota of each donor grouping. Radar plots display relative abundance of genes in metabolic pathways with significant^58^ mean decrease accuracies differentiating between indicated groups by random forest supervised machine learning, with the associated out- of-bag error indicated below each key for comparing donor A against the donors B & C. Values below each pathway label indicate the number of unique genes identified as belonging to each pathway from all contributing species.

To assess differences in microbial taxonomic composition among the donor metagenomes, we employed MetaPhlAn2^21^ using the default settings. Through this analysis we found a dominance of Bacillota (formerly Firmicutes) across all the donors, with Clostridiaceae being enriched relative to Ruminococcaceae in donor A samples (high-success FMT) compared to those from donors B & C (low-success FMT) (Fig. 1B). There also was relatively little difference in alpha-diversity between the grouping (Fig. S2A & S2B). Next, we sought to determine superorganism-level functional capacity. To accomplish this goal, we first performed a *de novo* assembly of contiguous genomic sequences (contigs) and then annotated gene content according to KEGG pathways. To measure the differential abundances of each pathway across donor groups we mapped metagenomic reads to annotated genes from each sample. Supervised machine learning was applied to these data to identify the functionalities that best differentiated the microbiomes. Comparing the donor A samples against donors B & C samples, it was revealed that glycosaminoglycan, fructose, mannose, arginine, and proline metabolism discriminated between donors and were significantly overrepresented in donor A (high-success FMT) samples (error rate: 8.33%) (Fig. 1C). Similarly, while achieving a lower overall accuracy, comparison of all successful outcome-associated samples against the FMT failure highlighted arginine & proline metabolism as well as glycosaminoglycan degradation as more abundant in communities associated with successful FMT (Fig. 1D). Since our *a priori* prediction was that metabolic interaction(s) with *C. difficile* are important for successful FMT, as a control, we performed the same analysis for non-metabolic pathway gene abundances but found substantially reduced ability to discriminate groups in either comparison (Fig. S2C & S2D). Ultimately the metabolic pathway abundances were more informative than analyses at the taxonomic level in defining traits of successful FMT thus supporting our hypothesis that FMT functions via metabolism rather than by the presence of specific species. However, the metabolic pathways identified in this analysis do not denote directionality for flow of metabolites leaving us unable to determine if identified pathways represent competition or cross-feeding with *C. difficile*.

To validate the observations from our single-site clinical study, we examined two publicly available datasets of 16S rRNA gene sequencing of samples across multiple donors in well-controlled, single-site studies of FMT efficacy^22,23^. Across both studies, successful clinical outcome was achieved in 79.8% of patients (71/89). Lachnospirales were largely the dominant taxa across donor samples (Fig. S1A) and was primarily composed of the genus *Blautia* (Fig. S1B), which has been associated with increased consumption of amino acids with additional probiotic properties^24^ and reduced abundance in CDI patient cohorts^25^. To explore possible differences in representation of metabolic pathways we assessed the imputed metagenomic content from ASV abundances using PICRUSt^26^. Results show a high abundance of genes involved in amino acid biosynthesis in successful FMT samples, supporting nutrient cross-feeding as a critical factor in rCDI resolution by FMT (Fig. S1C).

### MAG-directed analysis identifies specific metabolic pathways across diverse bacterial groups that associate successful FMT

Next, we sought to move past a superorganism view of metabolism by leveraging our metagenomic data to make a more discrete analysis of metabolic contributions of individual species to FMT efficacy. Contigs were clustered to yield putative Metagenome-assembled genomes (or MAGs) and curated for quality and potentially contaminating regions (Fig. S3). Gene content within each MAG was annotated with a protein-level 60% minimum identity and 1e-3 E-value to the KEGG GENES database, and putative species-level classifications were assigned to those MAGs with >50% single taxonomic origin for annotated genes. MAGs with the identical classification within single samples were collapsed and unique gene annotations were conserved to make the final collection of MAGs. Remaining MAGs were then validated for both contamination and completeness, which resulted in an overall reduction from 2411 draft MAGs to 463 high-quality validated MAGs with a median of 1876 annotated genes each for downstream analysis.

To focus specifically on association of MAG-encoded metabolism with FMT outcome, we then quantified metabolic pathway abundances within each MAG by mapping metagenomic reads from their associated samples and tabulating gene abundances for those KEGG pathways. Next, we employed supervised machine learning for these gene abundances to uncover pathways that best segregate MAGs associated with the highest success rate (Donor A). The resulting predictive model had a high overall accuracy of 74.1%, and those pathways with the highest predictive value associated with successful rCDI resolution involved multiple *C. difficile* carbon sources including genes associated with amino acid biosynthesis, as well as the metabolism of N-acetylglucosamine, sucrose, and galactose (Fig. 2A).

**Figure 2:**
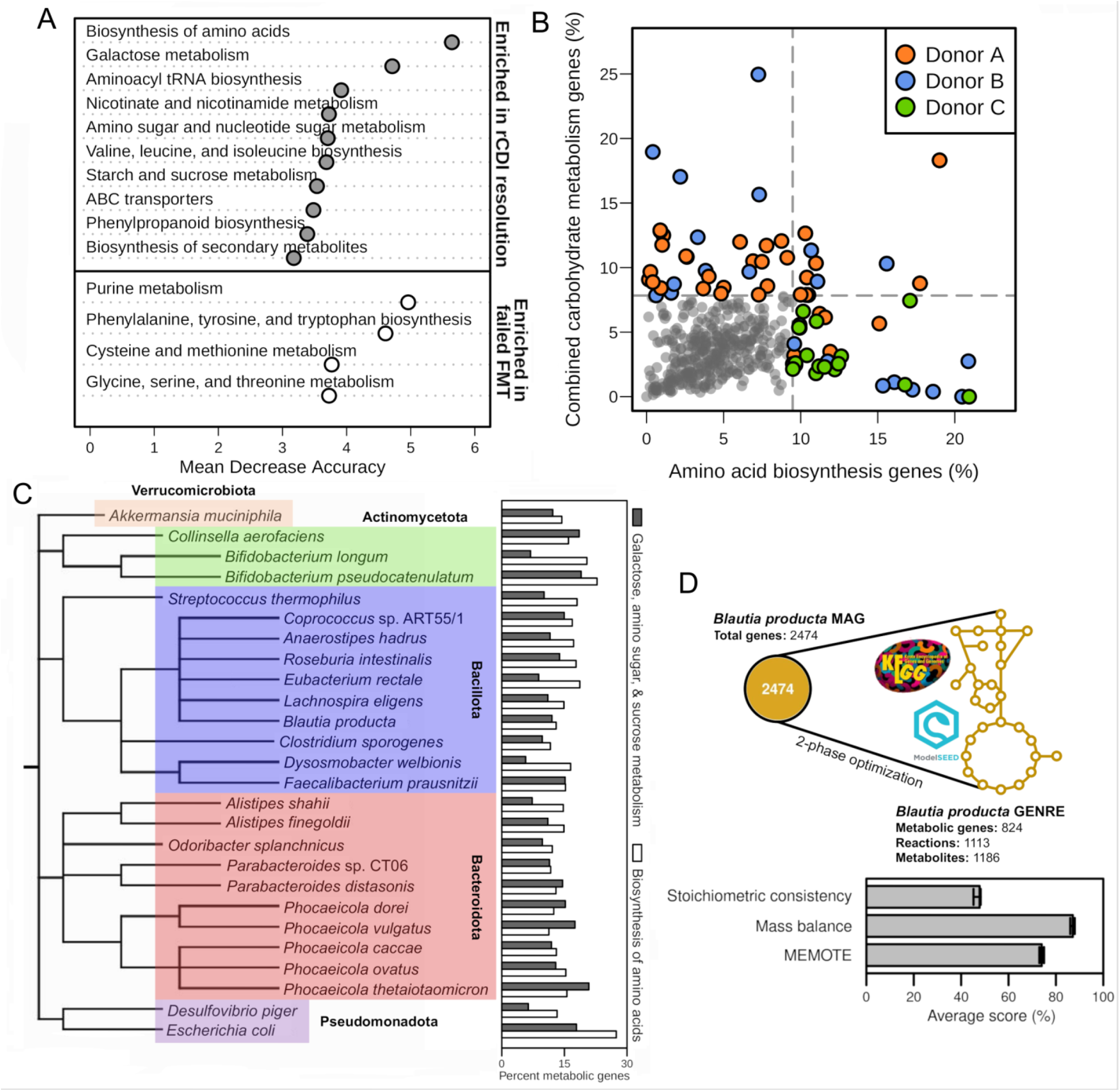
MAG-based analysis identified specific metabolism and contributing bacterial taxa that were associated with successful rCDI resolution. MAGs within each assembled metagenome were clustered and curated based on sequence composition and contig abundance. **(A)** Significant features^58^ from random forest supervised machine learning discriminating metabolic pathway gene abundances for MAGs associated with either successful or failed rCDI resolution. Groups indicate which FMT response was associated with the higher median gene abundance in each pathway. The resultant model had an OOB error of 25.88%, a PPV of 66.67%, and an NPV of 81.58%. **(B)** Scatterplot of MAG pathway abundance for amino acid biosynthesis and select carbohydrate metabolism (consolidation of galactose metabolism, amino sugar and nucleotide metabolism, and starch and sucrose metabolism), identified by machine learning analysis. Ratio of total genes in each MAG was calculated for each MAG, and a 90th quantile cutoff was applied to highlight 84 MAGs, which are colored by donor of origin. **(C)** Species-level classifications and phylogeny for the 26 GENREs used in downstream metabolic interaction simulations. To the left of the phylogeny are the percent of metabolic genes each pathway represents within the MAGs associated with each species. **(D)** Example *de novo* GENRE reconstruction from a curated MAG annotated as *Blautia producta*. The platform utilizes KEGG ortholog with associated ModelSEED metabolic reaction annotations to assemble scaffold models. Below the schematic are quality metrics for all 26 draft GENREs created for this study; Stoichiometric consistency, Reaction Mass Balance, and overall MEMOTE score.

Focusing on those pathways highlighted by the previous analysis, we sought to identify MAGs enriched for genes in amino acid biosynthesis, combined galactose metabolism, amino sugar & nucleotide sugar metabolism, and starch & sucrose metabolism pathways. We then calculated the fraction of genes within each MAG dedicated to each pathway and examined these ratios together to identify those MAGS with the highest capacities for either form of metabolism (Fig. 2B). To isolate the MAGs most associated with FMT success, we applied a strict abundance cutoff (>90th percentile) in either pathway category. This filter resulted in 85 MAGs of interest from across all three donors which contained the highest levels of either amino acid or carbohydrate metabolism, of which 63.53% were derived from successful FMT-associated samples (Table S4). Consolidation of MAGs with identical species annotations across samples then reduced this value to 44 MAGs (Table S2). Further integration of our metabolic analysis of MAGs with literature-based manual curation resulted in a limited collection of 26 MAGs that contained members across dominant phyla of the gut microbiome, possessed maximum metabolic association with successful FMT (Donor A), and had culturable isolates available (Fig. 2C).

### Simulated metabolic interactions with *C. difficile* revealed candidate probiotic bacterial species

To measure the directionality of each metabolic relationship with *C. difficile*, we sought to reconstruct the metabolism encoded in each of the 26 MAGs. While the gene and enzyme annotation level does not show directionality, flux balance analysis (FBA) based simulations allow for a clearer prediction of net consumption in *B. producta* and net production in *B. longum*. To do this, we developed a novel model building platform based on two slightly varied implementations of parsimonious flux analysis (Fig. 2D). Briefly, after mapping annotated genes to metabolic reactions, those gene-associated reactions were excluded from an overall flux minimization objective formulation to add the fewest reactions necessary to achieve growth (i.e., positive biomass reaction flux)^27^. Then, a minimum uptake rate for a common subset of minimal media components was applied during gapfilling, and the same process as the previous step was repeated, improving the ability of GENREs to grow in a greater variety of environmental conditions. We assessed the quality of the resultant GENREs using multiple quality metrics widely accepted within the field including stoichiometric consistency, overall mass balance of reactions, and MEMOTE^28^ score (a recently published tool for evaluating GENRE quality). This evaluation indicated that the GENREs produced here were high quality with a median MEMOTE score of 78% and flux/mass consistency in line with previous automated reconstructions.

To infer directionality of ecological interactions between the candidate probiotic strain GENREs and the C. *difficile* GENRE, we created a novel algorithm that characterizes potential edges of interaction as well as their degree of impact on growth during either cooperation or competition through separate iterative growth simulations. In each iteration, GENREs progressively increase levels of cooperation or competition for metabolites until no new interactions are found. The approach isolates the distinct metabolic impacts each microbe may have to reduce risk for false-negatives and maximize the ability to detect all possible edges of metabolic interaction(s). We performed simulations for all 26 MAGs generated in combination with the GENRE of the hypervirulent isolate *C. difficile* str. R20291 (iCdR703) and we found during cooperation simulations that both amino acids and carbohydrates were shared with *C. difficile*, with leucine, phenylalanine, and glucose-1- phosphate being the most frequent. Interestingly, there were relatively very few instances where *C. difficile* supplied metabolites to a partner species and mostly participated in largely one directional relationships (Fig. S4A). Competitive simulations identified almost exclusively amino acids, namely leucine and serine, as the edges of competition with other GENREs. Surprisingly, many of the bacteria identified in the analysis were more able to cross-feed rather than compete with *C. difficile*. While our initial hypothesis was that competition for nutrients was the most important interaction for FMT, these results support that metabolic cross-feeding may be a driving force behind successful FMT.

To validate our ecological interaction predictions, we selected two species with some of the highest predicted impacts on *C. difficile* growth *in silico* through either cooperation or competition for further testing. *Bifidobacterium longum* was a top cooperator and *Blautia producta* was one of the strongest competitors tested (Fig. 3B). This analysis revealed that the *B. longum* GENRE shared multiple preferred carbon sources with *C. difficile* including L-proline, ornithine, fumarate, and glucose-6-phosphate (Fig. 3C). Alternatively, *B. producta* was predicted to compete with *C. difficile* for exclusively amino acids, most strongly for asparagine, alanine, and threonine (Fig. 3D). To support these predictions *in vitro*, we employed a method of successive bacterial growth in conditioned rich culture medium that our group has previously utilized^29,30^ for measuring specific axes of metabolic interaction between species (Fig. 3E). Utilizing type strains *B. longum* ATCC 55813 and *B. producta* DSM 2950, we generated conditioned BHI + 10% FBS from either species through 24-hour incubation, followed by sterilization; 24-hour incubation ensures that species were grown to saturation, controlling for differences in kinetics and bacterial load. *C. difficile* str. R20291 was then grown for 18 hours in each conditioned medium to assess growth. As predicted by the *in silico* analysis, *C. difficile* grew poorly in *B. producta* conditioned medium which indicated a greater degree of nutrient depletion than in *B. longum* conditioned medium, which supported relatively high *C. difficile* growth (Fig. 3F).

**Figure 3:**
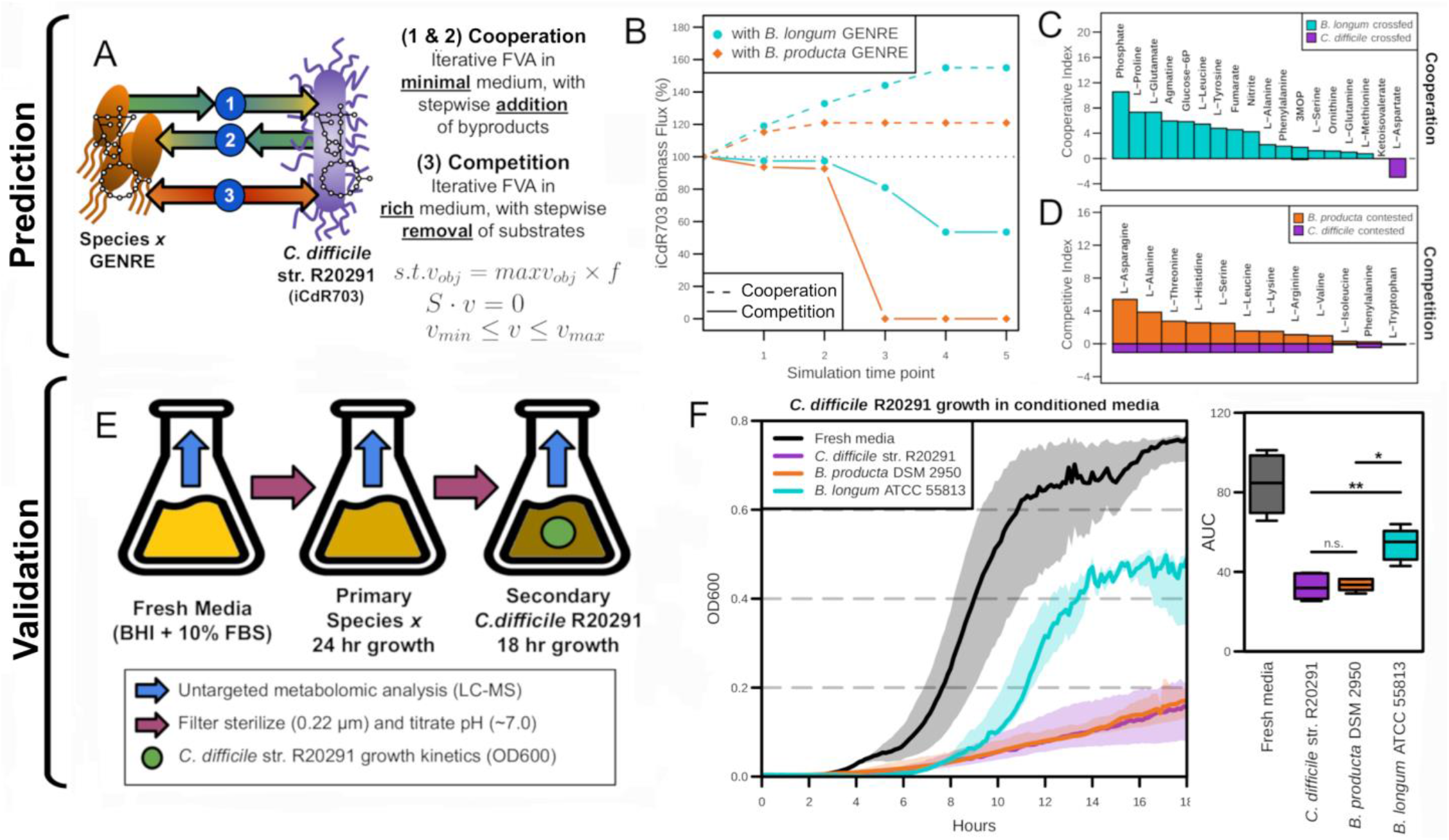
Prediction and validation of metabolic interactions between candidate probiotic strains and *C. difficile*. **(A)** Schematic and constraints for *in silico* simulation of metabolic interactions between probiotic and *C. difficile* GENREs. Simulations were initialized by calculating the combined minimal media components of both GENREs and setting flux ranges accordingly. **(B)** *C. difficile* str. R20291 GENRE (iCdR703) biomass flux (computational analog for bacterial growth) following each step of iterative growth and metabolic interaction simulations with interacting species. **(C)** Cooperative indices for predicted axes of cooperation between the *C. difficile* GENRE and the MAG-enabled GENRE of *Bifidobacterium longum*. *C. difficile* cooperative indices are represented as negative. **(D)** Competitive indices for predicted axes of metabolite competition between the *C. difficile* GENRE and the MAG-enabled GENRE of *Blautia producta*. *C. difficile* cooperative indices are represented as negative. **(E)** Schematic for *in vitro* validation of growth inhibition and metabolic crosstalk between candidate species and *C. difficile* str. R20291 through conditioned media experiments. Conditioned media was generated for each potential interactor species as well as *C. difficile* str. R20291 itself as a positive control of competition for growth substrates. **(F)** 18-hour growth of *C. difficile* str. R20291 in the indicated conditioned BHI (+10% FBS) medium (n=5). Area under the curve (AUC) was calculated in each replicate and significant differences determined by Wilcoxon rank-sum test with Benjamini-Hochberg correction.

To further validate ecological predictions, we next performed untargeted LC-MS profiling on aliquots of conditioned media from each step of the experimental procedure with *B. longum* and *B. producta* (Fig. 3E). Metabolites included in downstream analysis met one of the following criteria: scaled intensity for the given metabolite significantly decreased in both *C. difficile* and the other species’ individual cultures, implying competition for that metabolite, or the intensities were significantly increased in other species’ individual cultures and then significantly decreased following *C. difficile* growth in the conditioned medium suggesting cooperation. This multi-layered analysis showed that the *B. longum* added a substantial number of metabolites to the medium that were subsequently consumed by *C. difficile* versus metabolites that were competed for (Fig. 4A). Concordantly, *B. producta* had an opposite phenotype, competing for many more metabolites than it shared with *C. difficile* (Fig. 4B). Among the metabolites identified via this analysis were numerous Stickland fermentation substrates (indicated by green stars), with no carbohydrate concentration changes meeting the same strict criteria.

**Figure 4:**
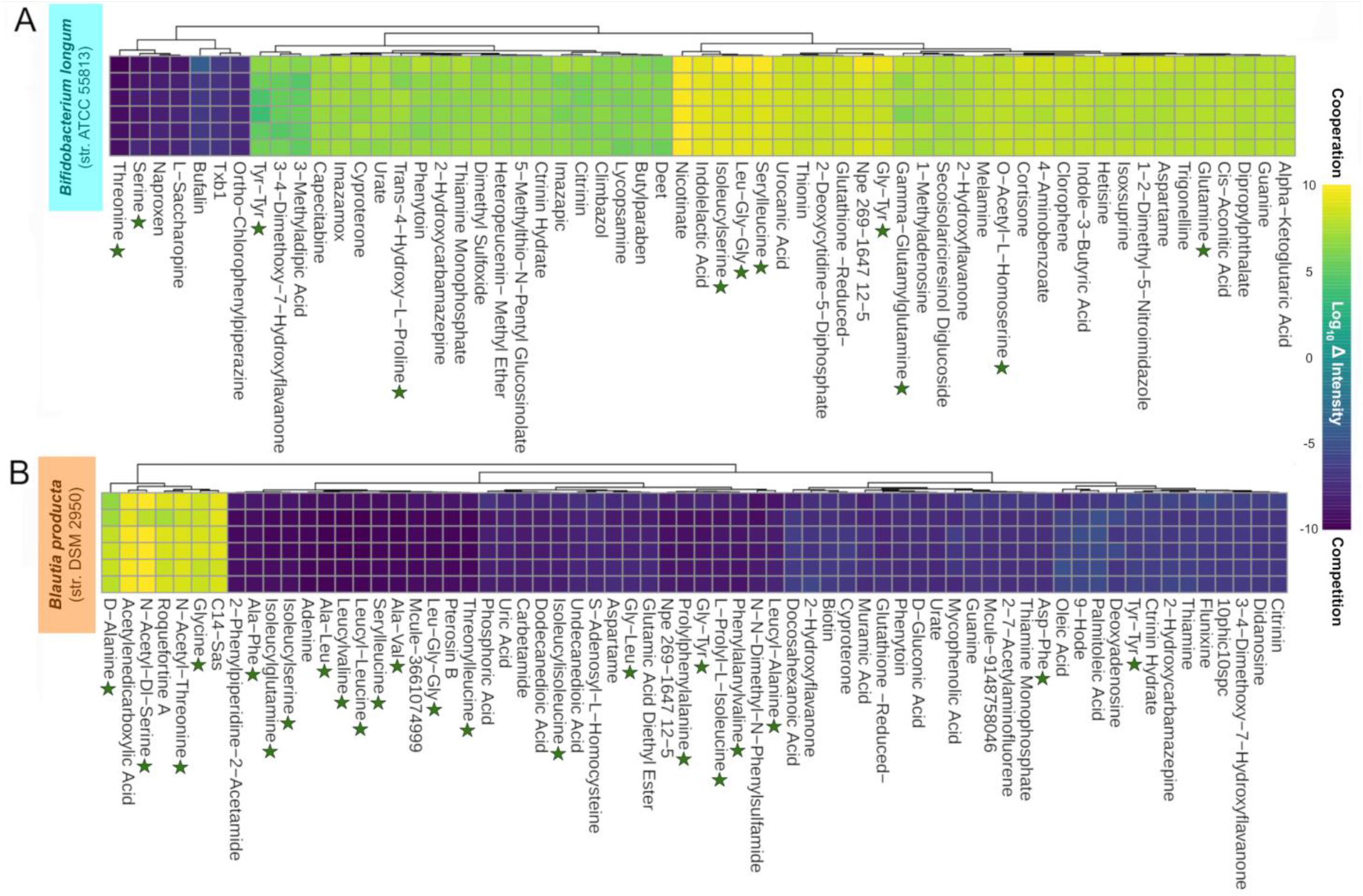
Metabolomics validation for ecological interactions with *C. difficile*. Untargeted LC-MS analysis from conditioned medium experiments at each labeled stage from Figure 3E for both validation species with *C. difficile* str. R20291 (n = 6 each); predicted cooperator **(A)** *Bifidobacterium longum* str. ATCC 55813 and predicted competitor **(B)** *Blautia producta* str. DSM 2950. Log_10_ Intensity refers to the transformed difference in relative intensity for metabolites between the medium before and after bacterial growth. For a change in relative scaled intensity of a given metabolite to be included in the analysis, it must be significantly decreased in both *C. difficile* and the other species’ individual cultures, or the intensities or the intensities must be significantly increased in other species’ individual cultures and then significantly decreased following *C. difficile* growth in the conditioned medium. Significant differences determined by Wilcoxon rank-sum test and green stars denote possible Stickland fermentation substrates.

### Cooperative consortium protects mice from CDI

Next, we assembled two distinct consortia that maximized either competition or cooperation with *C. difficile*. Membership of each consortium was chosen based on two criteria: first, species with the largest sum of competitive or cooperative indices, respectively, and second a unique range of individual metabolite interactions with *C. difficile* (Table S3). We limited each consortium to the top four species in each category to streamline laboratory cultivation of strains and promote reproducibility (Table 1); importantly, combined *in silico* and *in vitro* validation supported collective effects on *C. difficile* growth with no detectable direct deleterious effects (Fig. S4B, S4C, S6B, & S6C). To test the effect of our systems-assembled consortia on the course of CDI *in vivo*, we utilized a murine model of CDI (Fig. 5A). Mice were sensitized to infection with antibiotic pretreatment, then challenged with *C. difficile* str. R20291. On days one and two, following pathogen challenge, animals were orally gavaged with freshly mixed overnight (24 hr) cultures of the cooperative consortium, competitive consortium, or sterile growth medium as a control and then monitored for disease progression. In parallel, another set of identical groups of animals were utilized to also monitor changes in survival, and strikingly all animals that were treated with the cooperative consortium survived the infection, while all mice that received either the competitive consortium or control succumbed to the infection (Fig. 5B).

**Table 1:** Bacterial strains in each consortia predicted metabolic interactions with *C. difficile*. Strain designations along with the top specific metabolite axes of interaction with C. difficile, the total number of metabolites that the GENRE shares/competes for with iCdR703, as well as the sum of each interaction index for all metabolites in each category.

| Consortia | Member | Primary interaction (total interactions) | Σ Competitive Index | Σ Cooperative Index |
| --- | --- | --- | --- | --- |
| Competitive | *Bifidobacterium pseudocatenulatum* ATCC 27919 | L-Isoleucine (11) | 18.34 | 30.68 |
| Competitive | *Phocaeicola vulgatus* ATCC 8482 | L-Alanine (12) | 38.85 | 45.61 |
| Competitive | *Eubacterium rectale* ATCC 33656 | L-Tryptophan (5) | 28.49 | 53.18 |
| Competitive | *Blautia producta* DSM 2950 | L-Asparagine (12) | 26.42 | 60.93 |
| Cooperative | *Bifidobacterium longum* ATCC 55813 | L-Aspartate (19) | 18.72 | 84.73 |
| Cooperative | *Escherichia coli* K12 | Glycine, L-Proline (18) | 6.06 | 160.87 |
| Cooperative | *Roseburia intestinalis* DSM 14610 | L-Proline, L-Phenylalanine (20) | 18.69 | 137.49 |
| Cooperative | *Streptococcus thermophilus* LMD-9 | L-Glutamate (23) | 20.49 | 109.94 |

**Figure 5:**
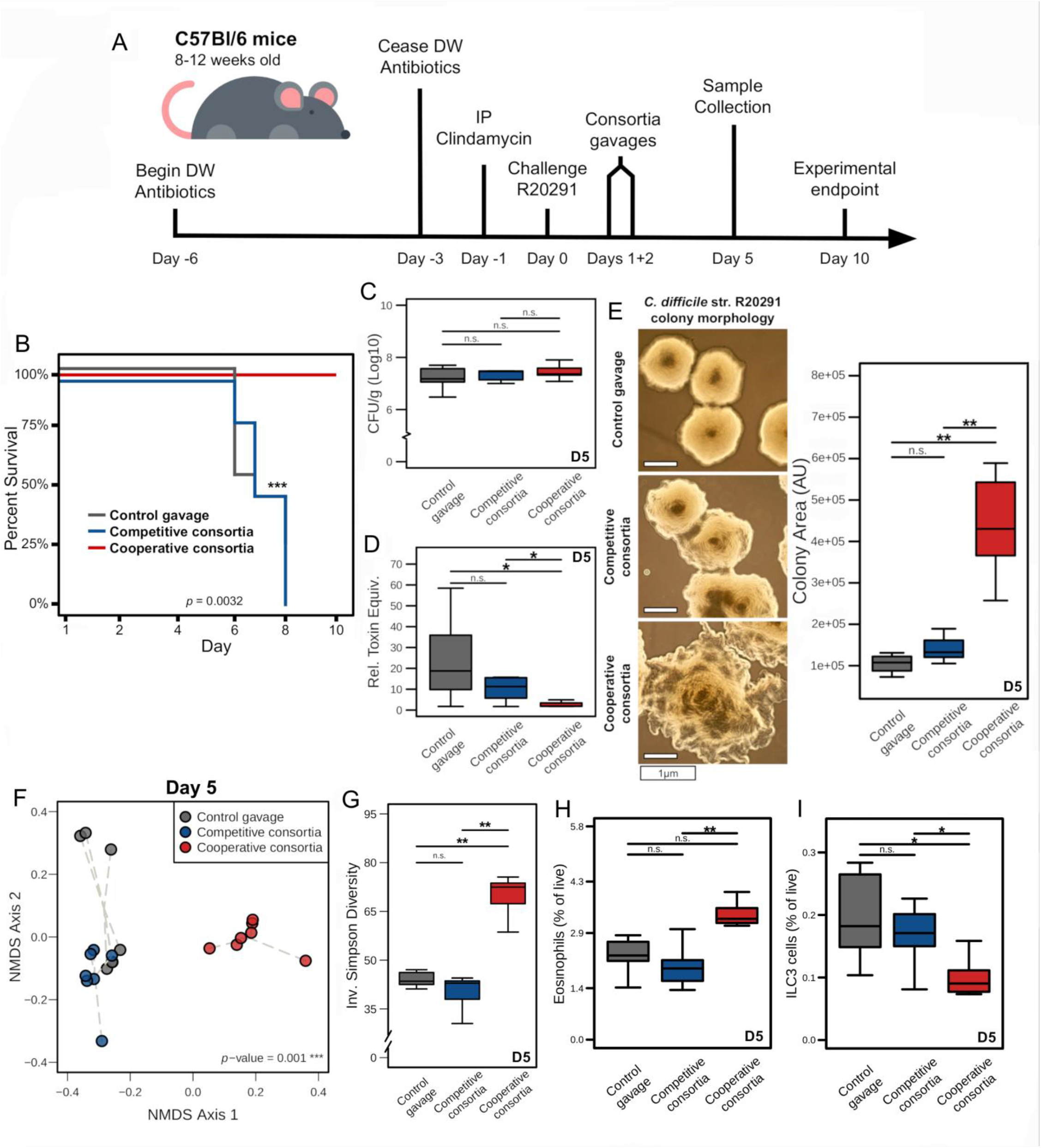
*in vivo* testing of targeted consortia in an animal model of acute CDI. **(A)** Experimental timeline of animal model used to test probiotic efficacy (n=14 per group). **(B)** Days post-infection survival curve for each experimental groups used. Significant difference determined by Kaplan-Meier test. **(C)** Total *C. difficile* str. R20291 CFU per gram of cecal content in mice (Day 5). Significant differences determined by Wilcoxon rank- sum test with Benjamini-Hochberg correction. **(D)** Relative concentration of *C. difficile* toxin per gram of cecal content in mice (Day 5). Significant differences again determined by Wilcoxon rank-sum test with Benjamini- Hochberg correction. **(E)** Microscopy of *C. difficile* colony morphologies from overnight growth on BHIS plated from cecal content of infected animals receiving the indicated probiotic consortia. Significant differences determined by Wilcoxon rank-sum test with Benjamini-Hochberg correction. **(F)** NMDS ordination of Bray-Curtis dissimilarities of cecal communities from each experimental group at the time of representative takedown (Day 5). Significant difference determined by PERMANOVA. **(G)** Inverse Simpson diversity of cecal communities from each experimental group (Day 5). Significant difference determined by Wilcoxon rank-sum test with Benjamini- Hochberg correction. Selected cell-type to live cell ratio from flow cytometric analysis; **(H)** Eosinophils or **(I)** ILC3 cells. Significant differences determined by Wilcoxon rank-sum test with Benjamini-Hochberg correction.

To assess protective factors directly or indirectly provided by the cooperative consortium, the experiment was repeated and animals were instead sacrificed on day five post infection to characterize pathogen burden, virulence expression and host-response. Based on *in vitro* data, we predicted that the consortia would impact *C. difficile* colonization, so we performed quantitative culture for total *C. difficile* colony forming units (CFU) from cecal content (Fig. 5C) as well as relative quantity of toxins A & B (Fig. 5D). This comparison revealed that there was no difference in the load of *C. difficile* across all groups; however, only the cooperative consortium was associated with significant decrease in detectable toxin (*p* = 0.012). As toxin expression has been previously linked to metabolism, we also measured individual colony (n = 6) area via microscopy image processing to assess possible distinct patterns of phase variation (Fig. 5E). This measurement revealed that colony morphology of *C. difficile* exposed to the cooperative consortium became significantly rougher than the other groups. Colony morphology is indicative of phase-variation in *C. difficile*^31,32^ that occurs through invertible elements upstream of the colony morphology regulator genes (*cmrRST*) and flagellar *flgB* operon which impact toxin regulation. We have previously demonstrated that this phenotypic heterogeneity is a metabolically sensitive phenotype^19^, and supports that the added consortia are interacting with the pathogen metabolically.

Based on the degree of change in CDI disease severity, we became interested in also measuring the collateral impacts of each consortium on the gut microbiota as well as host response to the infection ^33^. To accomplish these goals, we performed 16S rRNA gene sequencing from cecal content, which revealed a significant shift in community structure in only those animals receiving the cooperative consortia (Fig. 5F). We also found that alpha-diversity of the cooperative-associated communities significantly increased relative to either of the other groups (Fig. 5G). Importantly, only sequences for a fraction of each assembled consortia were detectable in any of the samples from the competitive (3/4) or cooperative-receiving (2/4) groups (Fig. S6A). To then measure indirect effects on FMT efficacy through the host^34^, we also characterized consortia-associated changes in immune response. Using spectra flow cytometry to characterize cells in the colonic lamina propria (Fig. S5A), we found that cooperative consortia promoted increased eosinophil levels (Fig. 5H) which has been associated with protection against CDI-associated mortality^9,35^. Additionally, significant decreases in ILC3 cells^36^ were observed in the same animals (Fig. 5I). These changes were also associated with no quantifiable differences in clinical scores between animals receiving the competitive consortia or control gavage (Fig. S5E).

## DISCUSSION

FMT has had extraordinary success in treatment of rCDI, with growing interest in leveraging this treatment for other diseases. In the current study, we developed a combined computational and experimental platform to identify potential probiotic bacterial species directly from human FMT metagenomic data. Leveraging rCDI as a proof of concept, we simulated ecological interactions for individual strains with *C. difficile* and identified a small collection of species with specific metabolic functions that reduced CDI-associated disease *in vivo*. This systems-level approach was demonstrably robust to a small metagenome sample size, and may have similar accuracy for other investigations of potential metabolic interactions between microbes in the future.

Our findings underscore the complex ecology inherent to FMTs and highlight the utility of targeting metabolism within the microbiota. While some amount of competition may be critical for ultimately eliminating *C. difficile* colonization, the success of FMT in reducing disease may be more contingent on metabolic cooperation. One line of evidence that may support this reasoning is that both *B. longum* and *E. coli* are prominent constituents of the infant microbiome where *C. difficile* is most often associated with asymptomatic colonization^37^. Our results support that amino acid cross-feeding of *C. difficile* by microbes present in FMT is likely a driving mechanism of reduced disease following infusion. While further mechanistic characterization of this pathway is needed, this interpretation agrees with previously appreciated negative genetic regulators of *C. difficile* toxin expression sensitive to intracellular levels of specific amino acids including L-proline^38^. Importantly, our results also suggest that FMT protects from CDI likely via a combination of effects on the pathogen, the microbiome, and the host response.

The methods and experimental designs of this pipeline were carefully considered to account for various challenges, such as sample size and prediction validations. Large and geographically diverse cohorts are often required to validate structural and functional features in microbiome data when the goal is to identify robust and consistent features. However, the goal of our pipeline was not to predict features that always occur in FMTs, but rather to select features that can successfully resolve CDI; this pipeline demonstrates the ability to use a small pool of FMT donations as a starting point for subsequent modeling, analyses, and experiments of potential therapeutic communities. We validated model predictions with an *in vitro* setting first which allows for tighter control of variables; importantly, the *in vitro* data agreed well with the model predictions (Fig. 3F, Fig. S4). It is important to emphasize that the models did not predict which of the communities would resolve disease but rather which communities had cooperative or competitive relationships with *C. difficile* in a highly controlled setting (as validated *in vitro*). We then tested the communities *in vivo* since such an environment is inherently less controlled. While engraftment of these introduced microbes into the host microbiome is not verified here, there is evidence that the therapeutic benefit of such communities does not require complete engraftment ^22,39^. Additionally, the therapeutic benefit of the cooperative consortium could arise from a subset of the four species in the consortium. Furthermore, as observed clinically, different strains of *C. difficile* can result in differences in disease; future work will need to explore how this pipeline can be applied to different *C. difficile* strains and to investigate what the minimal therapeutic consortium might be.

Systems biology approaches have been successfully applied to a variety of questions in *C. difficile* research. For example, a paired analysis of Environmental and Gene Regulatory Influence Networks (EGRIN) and metabolic models of *C. difficile* was used to identify the metabolic interactions between *C. difficile* and commensals at the microbial pathway and metabolite level^16^. Additionally, systems analysis of the intestinal microbiome is being leveraged to identify biomarkers that can successfully predict recurrence of CDI^40,41^. Furthermore, industry-based efforts have been made to create targeted bacterial consortia. Seres Therapeutics has designed an encapsulated version of a human FMT which can be delivered orally^15^ and Vedanta Biosciences is testing a bacterial consortia therapy that focuses on prevention of recurrence in high-risk patients (Clinical Trial ID: NCT03788434). However, to our knowledge, our approach is the first instance of a computationally-selected consortium using sequencing data from human FMT that has shown efficacy in an animal model of active infection and disease. Complete resolution of active CDI through non-invasive therapies will be a significant advancement within the field. Additionally, the systems biology pipeline created here can be used to identify additional tailored consortia against rCDI, as well as other collections of species that interface with host metabolism and related disease processes.

## MATERIALS & METHODS

### Clinical sample selection

The FMT donor preparations (OpenBiome; Boston, MA) used in this study were all part of the FMT cohort described previously (UVA IRB-HSR #18229)^20^. Two aliquots (1 mL each) were removed from each donor preparation used on the day of the patient’s FMT and stored at -80°C. Three donors were selected based on the number of donations and on patient outcome (i.e., FMT success/failure). A total of 119 patients with confirmed rCDI were treated with FMT and followed to determine treatment efficacy. A failed FMT was defined as recurrence of diarrhea with a positive stool ELISA for *C. difficile* toxin or positive *tcdB* PCR within three months of receiving FMT. From over the course of the study, four samples from each donor were chosen for DNA isolation and sequencing, controlling as much as possible for temporal effects within the donors as well as recipient-specific effects.

### DNA processing and high-throughput sequencing

Aliquots were thawed on ice, fecal material was centrifuged (14,000 rpm for 1 min), then resuspended in solution CD1 (800 µL) and processed according to the manufacturer’s instructions (QIAamp PowerFecal Pro DNA Kit). Total DNA was measured using the DeNovix Broad Range dsDNA kit on a DeNovix fluorometer and sent for paired-end 150bp shotgun sequencing using the Nextera XT kit standard protocol. Sequencing was performed on an Illumina HiSeq 2500. Libraries for V4 16S rRNA gene sequencing were generated using the established paired-end 250bp amplicon sequencing protocol from Kozich et. al. and sequenced on Illumina NextSeq 550. All sequencing was performed with GENEWIZ, Inc.

### Sequence read processing and analysis

Paired-end 16S rRNA gene sequencing reads from previously published clinical studies were obtained from the NCBI SRA (SRP064361 and SRP070464), and processed using the standard DADA2^42^ (v1.16) workflow (https://benjjneb.github.io/dada2/bigdata.html). Family and genus-level ASV assignment was performed using the SILVA SSU NR99 r138.1 database. Abundance analysis of ASVs was performed as described in the Statistical methods section below.

Raw metagenomic read quality trimming was performed using Sickle (>Q25), and contigs were assembled using Megahit^43^ with the following settings: minimum kmer size of 87, maximum kmer size of 127, and a kmer step size of 10. A minimum assembled contig size of 1500 bp was used for all downstream analysis. Full community putative gene content was identified with Prodigal^44^ and annotated with DIAMOND blastp^45^ against the KEGG^46^ protein database. Curated reads were mapped to genes with Bowtie2^47^, then PCR and sequencing errors were removed with PICARD MarkDuplicates to obtain final read abundances for “superorganism” level analyses within FMT subcommunities.

In the next phase, read abundances and computed tetranucleotide frequencies for each contig were used with Metabat2^48^ to cluster raw Metagenome-Assembled Genomes (MAGs). MAGs were decontaminated of possible erroneous contigs and pruned using CheckM^49^ (annotation kingdom or better, completeness ≥50% and contamination ≤33.3%) and excessively small MAGs were removed from downstream analysis (<1.3 Mb and <1354 putative genes; based on *Pelagibacter ubique* smallest free-living bacterium) to yield final high-quality MAGs. Gene content of each MAG was then called and annotated as in the previous section. Metagenomic reads were aligned to annotated genes of each MAG to assess relative abundance of gene/metabolic pathways and overall abundance of MAGs associated with positive outcome. Genus-level identification was performed by assessing for ≥40% annotated amino acid sequences in each MAG annotated as belonging to the same bacterial genus. Species identity of MAGs was determined with full length 16S rRNA gene BLAST and consensus gene annotation when possible. Full core functional content of final MAGs for downstream analysis was collectively validated through CheckM completeness-included annotations (Figure S3A).

### MAG-based *de novo* GENRE construction

KEGG + ModelSEED universal reaction database was adapted from a previous GENRE-building effort from our group^50^. Additional curation was performed to add new annotations (KEGG^46^ r88.1) ModelSEED Biochem DB (https://github.com/ModelSEED/ModelSEEDDatabase).

For each MAG, gene annotations were then mapped to corresponding metabolic reactions to create initial scaffolds for subsequent gapfilling. Gram +/- specific biomass formulations (Table S2) were then added to each GENRE scaffold based on taxonomic annotations from the previous analysis. Gapfilling was then achieved using the universal reaction database through a two-phase optimization approach:

Flux balance analysis (FBA) is a mathematical method for analyzing metabolic flux through an *in silico* metabolic network^51^. As in parsimonious-FBA, in the first phase each reversible reaction is separated into individual forward and reverse reactions *v_irrev_*. Linear coefficients *g* for each vector are now integrated (0 for reactions associated with a gene, and 1 for those that are not) and an objective function is then defined to minimize that total of flux across the entire model. These steps are all performed subject to a lower bound on the biomass flux (*v_obj_*) set to a fraction (*f*, set to ≥1%) of the previously identified maximum biomass flux (*max v_obj_*) and mass-balance constraints. Optimization (standard FBA) is performed and reactions found in the solution to have ≥1×10^5^ flux that are not in the initial scaffold GENRE are then added to the model.

In the second phase, additional constraints are added to the optimization where the input flux for select core growth substrate exchange reactions *v_e_* (Table S2) are required by setting a negative maximum flux (*k*, set to -0.01). As in the previous step, FBA is performed and new reactions found in the solution to have ≥1×10^5^ flux are added to the scaffold GENRE to yield the gapfilled model. Final GENREs in downstream analyses met size and quality minimum inclusion criteria of ≥180 genes and ≥400 reactions (sizes based on *E. coli* core model *i*AF126045), with <20% of metabolites participating in possible mass-generating loops^52^. MEMOTE^28^ was then run on each MAG-enabled GENRE resulting in a median quality score of 78% (Fig. 2F)^27^.

### Metabolic interaction predictions

First, each GENRE was constrained to achieve the minimum allowable optimal biomass of fraction *f* (≥25%) in complete media, at which point the minimal media components necessary for this growth rate were then calculated subject to standard mass balance and reaction flux constraints. Then for each model in the simulation (iCdR703 and interaction partner) the union of results from these calculations is then found and the extracellular exchange reactions of both models are set to mirror this combined list. Initial exchange flux ranges for the available metabolites in competition simulations were an *e_min_* of -1000 (lower bound) and an *e_max_* of +1000 (upper bound). In cooperative simulations initial *e_min_* was -1000. Flux variability analysis (FVA) was then performed for both GENREs focusing only on extracellular exchange reaction ranges. For those active exchange reactions for metabolites present in both GENREs, the lower or upper bound was modified by the given coefficient *c* (0.5) if calculating competition or *s* (100) if calculating cooperation.

During either type of interaction simulation, the opposite directions’ coefficient was set to 1 or 0 (*c* or *s*). Joint FVA was then repeated for the newly constrained GENREs until biomass flux either ceased to change or no new edges of interaction were found. Cooperative index for each metabolite was calculated as the exported metabolite exchange flux divided by the recipient GENRE’s maximum biomass flux. Alternatively, the competitive index for metabolites was calculated as the given metabolite’s absolute exchange flux divided by the importing GENRE’s maximum biomass flux, multiplied by the number of iterations that metabolite was contested.

### Strain acquisition, cultivation, and identification

Lyophilized type strain isolates for taxa identified through commercial vendors ATCC (https://www.atcc.org) or DSMZ (https://www.dsmz.de/) as noted in the individual species identifiers. All strains were reconstituted and grown anaerobically (6% H, 20% CO_2_, 74% N_2_) at 37°C in liquid Brain Heart Infusion medium (BHI; BD Bacto Ref:237500 Lot:1194923) supplemented with 10% Fetal Bovine Serum (FBS; Gibco Ref:A38403-02 Lot:2240781RP), and streaked on solid agar of the same medium (1.5% agar). Sanger sequencing of full 16S rRNA gene (Genewiz Inc., 27F - 1492R primers) was performed from all cultivated isolates prior to storage and experimentation to ensure correct isolates.

### Conditioned medium experiments

Successive conditioned media experiments were performed as described previously^29,30^. Single colonies from each potential consortium member growing on BHI + 10% FBS solid medium were inoculated to fresh liquid BHI + 10% FBS and conditioned media from each strain was prepared by growing each species for 24 hr anaerobically at 37 °C. The resulting culture was centrifuged at 3500 rpm for 10 min and the supernatant was returned to pH 7.0 using 1M NaOH then filter sterilized (PVDF membranes with 0.22 μm pore size). Aliquots of spent medium were stored at −80 °C, and individual aliquots were thawed and equilibrated in the anaerobic chamber overnight prior to *C. difficile* inoculation. Similarly, *C. difficile* single colonies were picked from overnight growth on BHI + 10% FBS solid agar and then incubated in liquid BHI + 10% FBS for an additional 24 hr. Prior to growth measurements, liquid overnights were back-diluted 1:3 into fresh liquid BHI + 10% FBS and incubated for 2 hr to initiate active growth. Conditioned media (250 μl each) was then inoculated with back-dilutions at a ratio of 1:100 in a 96 well plate (Corning Ref:3596), and OD_600_ was measured every 5 min for 18 hr (Tecan Infinite M200 Pro).

For conditioned medium supplementation experiments, sterile 24 hr growth conditioned medium from *C. difficile* str. R20291 was diluted 1:1 with sterile aliquots of either cooperative or competitive consortia-depleted media (24 hr growth). Positive and negative controls of fresh BHI + 10% FBS or additional *C. difficile*-depleted media were performed simultaneously. Growth (OD600) was then measured as in the previous conditioned media experiments and AUCs were calculated from the resultant curves.

### Disk diffusion assay

The entire surface of standard anaerobic BHIS agar plates was inoculated and spread with 100 µL of overnight *C. difficile* str. R20291 growth. Plates were air-dried for 10 minutes, at which point diffusion disks (Sigma #741146) saturated with 25 µL of liquid were added to the surface with replicates for each group. Plates were incubated at 37C for 24 hr and inhibition zone diameter was measured. Vancomycin concentration and area measurements were performed according to the guidelines established by the National Committee for Clinical Laboratory Standards. Growth inhibition diameter measurements from disk diffusion assay for *C. difficile* str. R20291 with the indicated treatments (n=6).

### Untargeted Mass Spectrometry

Liquid chromatography-high resolution mass spectrometry (LC-HRMS) analysis from conditioned media was performed in the UVA Metabolomics Core as previously described^53^. In summary, randomized sample injection was performed via a Thermo Vanquish UHPLC, and separation of the metabolites was achieved using Thermo Accucore C18 column (Thermo Scientific; 2.1 by 100 mm, 1.5 μm). For the 15-minute gradient, the mobile phase consists of solvent A (0.1% formic acid in water) and solvent B (0.1% formic acid in methanol). The gradient was as follows: 0 to 8.0 min 50% solvent B, 8.0 to 13.0 min held at 98% solvent B, and 13.1 to 15.0 min revert to 0% solvent B to re-equilibrate for next injection. Spectra were acquired on Thermo ID-X Tribrid MS, using both positive and negative mode. Data was acquired in full MS mode (1 μscan) at a resolution of 120,000 with a scan range of 67 to 1,000 m/z at a normalized automatic gain control target of 25%, and 50 ms of maximum injection time was allowed. Fragment masses were detected by Orbitrap at a resolution of 30,000. Raw spectra were analyzed in Compound Discoverer 3.1 with standard settings. Compound annotations were performed by searching the ddMS2 masses in mzCloud, ChemSpider, KEGG database, and an in-house database of IROA Mass Spectrometry Metabolite Library of Standards (https://iroatech.com/).

### Animal infection model

Murine *C. difficile* infections were carried out as previously described^9^. Five- week-old C57BL6 male mice were purchased from Jackson Laboratories. All mice were housed under specific pathogen-free conditions at the AALAC accredited University of Virginia animal facility. Experiments were performed according to the provisions of the Animal Welfare Act and were approved by the University of Virginia Institutional Animal Care and Use Committee. Mice were maintained on the Envigo control diet (TD.05230) for three weeks before treatment with antibiotics and throughout the infection model. Prior to beginning the experimental model, mice from across litters were randomly assigned to respective groups. To make mice susceptible to infection, mice received an oral antibiotic cocktail containing 45 mg/L vancomycin (Mylan), 35 mg/L colistin (Sigma), 35 mg/L gentamicin (Sigma) and 215 mg/L metronidazole (Hospira) *ad libitum* for 3 days. The animals were switched back to plain drinking water two days before the infection. The day before infection, mice were injected intraperitoneally with 0.016 mg/g of clindamycin. The next day, mice were orally challenged with 10^3^ *C. difficile* str. R20291 spores in 100 µl of sterile water via oral gavage. During the infection, mice were monitored daily for weight loss, as well as clinical signs of disease. Clinical score criteria included weight loss, coat condition, activity level, diarrhea, posture and eye condition for a cumulative clinical score between 1 and 20. Weight loss and activity levels were scored between 0 and 4 with four being greater than or equal to 25% loss in weight. Coat condition, diarrhea, posture, and eye condition were scored between 0 and 3. Diarrhea scores were 1 for soft or yellow stool, 2 for wet tail, and 3 for liquid or no stool. Mice were euthanized if severe illness developed based on a clinical score ≥ 14. On days two and three post *C. difficile* challenge, 100 µl of each consortium, prepared from 24-hour cultures, was administered to mice via oral gavage. A third group received vehicle control.

### In vivo C. difficile CFU quantification

Cecal content samples were weighed and serially diluted under anaerobic conditions (6% H, 20% CO_2_, 74% N_2_) with anaerobic PBS. Plating for total *C. difficile* str. R20291 CFU was performed by plating diluted samples on BHI plates plus taurocholate (10%), cycloserine (0.5%), and cefoxitin (0.5%) at 37°C for 24 hr under anaerobic conditions after which, the resultant colonies were counted across all experimental replicates.

### Relative toxin quantification

Supernatants were tested for the combined presence of Toxin A/B using the ToxA/B II kit (TechLab) according to manufacturer’s directions with the following modifications. Briefly, supernatants from dilutions of cecal contents used for *C. difficile* R20291 CFU quantification were directly added to assay wells. The goal was to measure the relative amount of toxins present in the sample compared to the positive control provided in the kit. To do this, increasing volumes of the positive control were used to construct a standard curve with the OD450 readout of samples calculated as relative toxin equivalents per gram of cecal content and normalized by the absolute quantity of *C. difficile* per gram of cecal content. We note that PCR can result in overcalls of toxin positivity^54^ and so our results are normalized to absolute quantity of *C. difficile* per gram of cecal content.

### Colony Microscopy

At least one colony from each animal was included in each comparison for *C. difficile* grown on BHIS from cecal content of infected mice. Microscopy images were taken on an EVOS XL Core cell imaging system at x4 magnification. Colony dimensions were determined using ImageJ (https://imagej.nih.gov/ij/).

### Immune Profiling

To isolate the lamina propria, colons were removed, longitudinally cut, and rinsed in cold Hank’s Balanced salt solution (HBSS) supplemented with 5% FBS and 25mM HEPES. Epithelial cells were removed by incubating each colon in pre-warmed HBSS supplemented with 15 mM HEPES, 5 mM EDTA, 10% FBS and 1mM dithiothreitol at 37C for 40 minutes in a shaking incubator at 220 rpm. The colon was then cut into smaller sections and digested in RPMI containing 0.17 mg/mL liberase TL (Sigma) and 30 mg/mL DNase (Sigma) at 37C in a shaking incubator. Following digestion, the tissue was passed through 40 and 100 uM cell strainers respectively, the cells were counted and re-suspended in FACS buffer (PBS with 2% FBS). Approximately 1 x 106 cells were plated in a 96-well plate and stained for flow cytometry. First, cells were Fc blocked (anti-mouse CD16/32 TruStain, BioLegend Cat #101320) and then stained with a fixable viability dye (Zombie NIR, BioLegend Cat #423105). The following antibodies were used for staining, CD19-PerCP-Cy5.5 ( BioLegend, Cat #115534), CD3-PerCP-Cy5.5 (BioLegend, Cat #100218), CD5-PerCP-Cy5.5 (BioLegend, Cat #100624), FcER1a- PerCP-Cy5.5 (BioLegend, Cat #1343420), Ly6C-AlexaFluor488 (BioLegend, Cat #128022), CXCR2- BV750 (BD, Cat #747816), Ly6G-BV650 (BioLegend, Cat #127641), SiglecF-AlexaFluor700 (ThermoFisher, Cat #56-1702-82), CD45-SparkViolet538 (BioLegend, Cat# 103179), CD8a-NovaFluorBlue610-70S (ThermoFisher, Cat#M003T02B06), CD4-APC Fire750(BioLegend, Cat# 100460), CX3CR1-Alexa Fluor 647(BioLegend, Cat# 149004), CD90.2- BV785 (BioLegend, Cat# 105331), CD11c- PE-Cy7 (BioLegend, Cat# 117318), CD11b- BV480 (BD, 566117), TCR Beta- BV570 (BioLegend, Cat# 109231), CD64- BV421 (BioLegend, Cat# 1393090), MHCII-BV605 (BioLegend, , Cat# 107620), CD127- PE-Cy5 (BioLegend, Cat# 1350160), CD86- PE-Dazzle 594 (BioLegend, Cat# 105042), CD206-PE (BioLegend, Cat# 141706), GATA-3- BV711(BD, Cat# 565449), ROR gammaT- APC (ThermoFisher, Cat# 17-6981-82), FoxP3- PerCP-eFluor710 (ThermoFisher, Cat# 46-5773-82). For intracellular transcription staining, the Foxp3/ Transcription Factor Staining Buffer Set (eBioscience, Cat #00- 5523-00) was used according to the manufacturer’s instructions. Flow cytometry data was acquired using the Cytek Aurora Borealis outfitted with 5 lasers including blue, violet, yellow-green, red and UV. Data analysis was performed using OMIQ.

### Statistical methods

All statistical analysis was performed in R v4.1.3. Rarefaction (11,000 reads per sample), nonmetric multidimensional scaling of Bray-Curtis dissimilarity, and permutational multivariate analysis of variance (PERMANOVA) analyses were accomplished using the vegan R package^55^. Significant differences for growth curve AUC, metabolite concentrations, *C. difficile* CFU, community diversity, relative toxin equivalents, colony morphology, and immune cell population counts were determined by Wilcoxon signed-rank test and Benjamini-Hochberg correction when necessary. Supervised machine learning was accomplished with the base implementation of Random Forest also in R^56^. Dissimilarity between *C. difficile* str. R20291 growth curves in consortium-conditioned media were determined using Dynamic Time Warping^57^.

#### Data and code availability

The GitHub repository for this study, with all programmatic code and GENREs described here, can be found at https://github.com/csbl/Jenior_FMT_2022. 16S rRNA gene sequencing reads were downloaded in raw FASTQ format from the NCBI Sequence Read Archive (SRP064361 and SRP070464). Screened metagenomic sequencing reads generated for this study have been uploaded to the NCBI SRA (PRJNA844211).

## FIGURE & TABLE LEGENDS

**Figure S1: Taxonomic composition and predicted functional capacity of microbiota in successful FMT donor samples.** Results from analysis of 16S rRNA gene sequencing collected from capsule donor samples with 79.8% (71/89) efficacy used in the previous study from Staley et. al.^22^. To avoid skewing results toward individuals with larger numbers of utilized capsules, a maximum of 5 samples were randomly selected (when possible) from each donor for analysis across 10 unique donor pools, resulting in 76 donor samples for downstream analysis. **(A)** Relative abundances of Family-level taxonomic representation from ASVs associated with successful resolution of rCDI. **(B)** Genus-level breakdown of the dominant Family (Lachnospirales) from panel A. **(C)** Median and IQR for gene abundances from top 25 most abundant pathways in imputed metagenomic content using PICRUSt. Median points are colored based on the pathway’s relationship to metabolism. **(D)** Breakdown of most abundant individual gene inferred abundances.

**Figure S2: Supplementary metagenomic analysis of FMT donor samples.** Comparisons of Inv. Simpson diversity from community structures inferred by MetaPhlAn2 analysis in **(A)** high success-associated versus low success-associated samples or **(B)** successful rCDI resolution versus failure. Differences calculated by Wilcoxon ranked-sum test. **(C & D)** Similar analysis to Figures 1C and 1D, instead limiting the predictive model to only non-metabolic pathway abundances at Superorganism-level.

**Figure S3: MAG quality filtering results.** Raw MAGs generated from unsupervised Metabat2 contig clustering were pruned first by a minimum size of ≥1.3 Mb and ≥1345 putative genes (based on *Pelagibacter ubique*, smallest genome of a free-living bacterium). CheckM analysis was then performed on both size-filtered MAG groups, and another filter was applied of >50% completeness and <33% contamination (defined within CheckM). CheckM quality metrics for the pre-filter and resultant post-filter MAGs are shown in **(A)** completeness and **(B)** contamination. Significant differences determined by Wilcoxon rank-sum test.

**Figure S4: Computational and *in vitro* validation of consortia interactions with *C. difficile*. (A)** Quantification of directionality for shared metabolites across all cooperative simulations performed with iCdR703. **(B)** *in silico* consortia impact on iCdR703 biomass flux. Interaction metabolites for each group were determined using the union of edges of interaction returned by individual simulations for component species. Lower bounds of exchange reactions for edges of potential competition were set to 0, and those for edges of potential cooperation were set to -1000. **(C)** Combined *in vitro* consortia effects on 18 hr growth of *C. difficile* str. R20291 at 37℃ in the indicated pre-conditioned media compared to fresh BHI + 10% FBS (n=5). Significant differences determined by Dynamic Time Warping and PERMANOVA.

**Figure S5: Additional immunological characterization of murine infection models with consortium gavage. (A)** Spectral flow cytometry gating strategy. Resultant densities of cell types of interest are as follows **(B)** Select innate immunity cell lineages with previous links the CDI response, **(C)** select adaptive immunity cell lineages with previous links the CDI response, and **(D)** additional cell lineages with large-scale changes between groups. All values reflect percentages of total live cells detected, and significant differences determined by Wilcoxon rank-sum test. **(E)** Clinical scores of mice following infection and bacteriotherapy (n = 9). Groups with no remaining animals are denoted by NA. Scores were based on weight loss, coat conditions, activity level, diarrhea, posture and eye condition for a cumulative clinical score between 1 and 20. Weight loss and activity levels were scored between 0 and 4 with four being greater than or equal to 25% loss in weight. Coat condition, diarrhea, posture, and eye condition were scored between 0 and 3. Diarrhea scores were 1 for soft or yellow stool, 2 for wet tail, and 3 for liquid or no stool. Mice were euthanized if severe illness developed based on a clinical score ≥ 14.

**Figure S6: Additional testing for consortia detectability and growth suppression from conditioned media. (A)** Detectability of 16S rRNA gene sequencing reads of the respective consortium members. **(B)** Growth inhibition diameter measurements from disk diffusion assay for *C. difficile* str. R20291 with the indicated treatments (n = 6). **(C)** Area under the curve results from 18-hr *C. difficile* str. R20291 growth in supplemented (50:50) liquid BHIS previously depleted by str. R20291 over 24 hours with the indicated growth medium variation. Significant differences were determined by Wilcoxon rank-sum test with Benjamini-Hochberg correction (n = 4).

**Table S1: Metagenomic sequencing read quality and assembly statistics**

**Table S2: All GENRE statistics and quality metrics**

**Table S3: Metabolic interaction simulation results for all GENREs**

**Table S4: MAG pathway abundance screening results**

## ACKNOWLEDGMENTS

The authors would like to thank Nicole Koropatkin for the donation of several bacterial strains used in this study. We also thank Nishikant Wase at the UVA Health System Metabolomics Core for support in the *in vitro* study design and mass spectrometry data generation. Additionally, Toxin A/B II detection kits were generously provided by James Boone at TechLab. Finally, the authors are thankful for the support of Alexander Jenior, who was instrumental in the initial drafting of the manuscript.

## FUNDING

This work was supported by funding from the U.S. National Institutes of Health awards R01AT010253 to J.A.P., R01AI124214 to W.A.P., F32DK124048 to J.L.L., and a pilot grant from the UVA Trans-University Microbiome Initiative to M.L.J. The funding agency had no role in study design, data collection/analysis, or preparation of the manuscript.

## AUTHOR CONTRIBUTIONS

M.L.J.—Conceptualization. Sample collection. Data generation and analysis. Drafting manuscript.

J.L.L.—Conceptualization. Sample collection. Data generation and analysis. Drafting manuscript.

G.L.K.—Sample collection. Clinical data gathering. Data generation. Editing manuscript.

L.A.P.—Clinical study execution. Sample collection. Editing manuscript.

D.A.P.—Editing manuscript.

W.A.P.—Supervision. Editing manuscript.

J.A.P.—Funding acquisition. Supervision. Drafting and editing manuscript.

## Notes

### Competing Interest Statement

The authors have declared no competing interest.

## REFERENCES

1. Shreiner, A. B., Kao, J. Y. & Young, V. B. The gut microbiome in health and in disease. Curr Opin Gastroenterol 31, 69–75 (2015).

2. D’Haens, G. R. & Jobin, C. Fecal Microbial Transplantation for Diseases Beyond Recurrent Clostridium Difficile Infection. Gastroenterology 157, 624–636 (2019).

3. Wang, S. et al. Systematic Review: Adverse Events of Fecal Microbiota Transplantation. PLoS One 11, e0161174 (2016).

4. Lessa, F. C., Winston, L. G., McDonald, L. C., & Emerging Infections Program C. difficile Surveillance Team. Burden of Clostridium difficile infection in the United States. N Engl J Med 372, 2369–2370 (2015).

5. Figueroa Castro, C. E. & Munoz-Price, L. S. Advances in Infection Control for Clostridioides (Formerly Clostridium) difficile Infection. Curr Treat Options Infect Dis 11, 12–22 (2019).

6. Kärki, T., Plachouras, D., Cassini, A. & Suetens, C. Burden of healthcare-associated infections in European acute care hospitals. Wien Med Wochenschr 169, 3–5 (2019).

7. Valdes, A. M., Walter, J., Segal, E. & Spector, T. D. Role of the gut microbiota in nutrition and health. BMJ 361, k2179 (2018).

8. Theriot, C. M. et al. Antibiotic-induced shifts in the mouse gut microbiome and metabolome increase susceptibility to Clostridium difficile infection. Nat Commun 5, 3114 (2014).

9. Frisbee, A. L. et al. IL-33 drives group 2 innate lymphoid cell-mediated protection during Clostridium difficile infection. Nat Commun 10, 2712 (2019).

10. Bakken, J. S. Fecal bacteriotherapy for recurrent Clostridium difficile infection. Anaerobe 15, 285–289 (2009).

11. Baktash, A. et al. Mechanistic Insights in the Success of Fecal Microbiota Transplants for the Treatment of Clostridium difficile Infections. Front Microbiol 9, 1242 (2018).

12. Wilson, K. H. & Perini, F. Role of competition for nutrients in suppression of Clostridium difficile by the colonic microflora. Infect Immun 56, 2610–2614 (1988).

13. Collins, J., Danhof, H. & Britton, R. A. The role of trehalose in the global spread of epidemic Clostridium difficile. Gut Microbes 10, 204–209 (2019).

14. Neumann-Schaal, M., Jahn, D. & Schmidt-Hohagen, K. Metabolism the Difficile Way: The Key to the Success of the Pathogen Clostridioides difficile. Front. Microbiol. 10, 219 (2019).

15. Khanna, S. et al. A Novel Microbiome Therapeutic Increases Gut Microbial Diversity and Prevents Recurrent *Clostridium difficile* Infection. J Infect Dis. 214, 173–181 (2016).

16. Girinathan, B. P. et al. In vivo commensal control of Clostridioides difficile virulence. Cell Host Microbe 29, 1693–1708.e7 (2021).

17. Bauer, E. & Thiele, I. From Network Analysis to Functional Metabolic Modeling of the Human Gut Microbiota. mSystems 3, e00209–17 (2018).

18. Bauer, E. & Thiele, I. From metagenomic data to personalized in silico microbiotas: predicting dietary supplements for Crohn’s disease. NPJ Syst Biol Appl 4, 27 (2018).

19. Jenior, M. L., et al. Novel Drivers of Virulence in Clostridioides difficile Identified via Context-Specific Metabolic Network Analysis. mSystems 6, e0091921 (2021).

20. Shin, J. H. et al. Outcomes of a Multidisciplinary Clinic in Evaluating Recurrent Clostridioides difficile Infection Patients for Fecal Microbiota Transplant: A Retrospective Cohort Analysis. J Clin Med 8, 1036 (2019).

21. Truong, D. T. et al. MetaPhlAn2 for enhanced metagenomic taxonomic profiling. Nat Methods 12, 902– 903 (2015).

22. Staley, C. et al. Successful Resolution of Recurrent Clostridium difficile Infection using Freeze-Dried, Encapsulated Fecal Microbiota; Pragmatic Cohort Study. Am J Gastroenterol 112, 940–947 (2017).

23. Staley, C. et al. Predicting recurrence of Clostridium difficile infection following encapsulated fecal microbiota transplantation. Microbiome 6, 166 (2018).

24. Liu, X. et al. Blautia-a new functional genus with potential probiotic properties? Gut Microbes 13, 1–21 (2021).

25. Daquigan, N., Seekatz, A. M., Greathouse, K. L., Young, V. B. & White, J. R. High-resolution profiling of the gut microbiome reveals the extent of Clostridium difficile burden. NPJ Biofilms Microbiomes 3, 35 (2017).

26. Langille, M. G. I. et al. Predictive functional profiling of microbial communities using 16S rRNA marker gene sequences. Nat Biotechnol 31, 814–821 (2013).

27. Jenior, M. L., Glass, E. M. & Papin, J. A. Reconstructor: A COBRApy compatible tool for automated genome-scale metabolic network reconstruction with parsimonious flux-based gap-filling. http://biorxiv.org/lookup/doi/10.1101/2022.09.17.508371(2022) doi:10.1101/2022.09.17.508371.

28. Lieven, C. et al. MEMOTE for standardized genome-scale metabolic model testing. Nat Biotechnol 38, 272–276 (2020).

29. Biggs, M. B. et al. Systems-level metabolism of the altered Schaedler flora, a complete gut microbiota. ISME J 11, 426–438 (2017).

30. Medlock, G. L. et al. Inferring Metabolic Mechanisms of Interaction within a Defined Gut Microbiota. Cell Syst 7, 245–257.e7 (2018).

31. Anjuwon-Foster, B. R. & Tamayo, R. A genetic switch controls the production of flagella and toxins in Clostridium difficile. PLoS Genet 13, e1006701 (2017).

32. Anjuwon-Foster, B. R. & Tamayo, R. Phase variation of Clostridium difficile virulence factors. Gut Microbes 9, 76–83 (2018).

33. Littmann, E. R. et al. Host immunity modulates the efficacy of microbiota transplantation for treatment of Clostridioides difficile infection. Nat Commun 12, 755 (2021).

34. Jan, N., et al. Fecal Microbiota Transplantation Increases Colonic IL-25 and Dampens Tissue Inflammation in Patients with Recurrent Clostridioides difficile. mSphere 6, e0066921 (2021).

35. Buonomo, E. L. et al. Microbiota-Regulated IL-25 Increases Eosinophil Number to Provide Protection during Clostridium difficile Infection. Cell Reports 16, 432–443 (2016).

36. Fachi, J. L. et al. Acetate coordinates neutrophil and ILC3 responses against C. difficile through FFAR2. J Exp Med 217, jem.20190489 (2020).

37. Penders, J. et al. Quantification of Bifidobacterium spp., Escherichia coli and Clostridium difficile in faecal samples of breast-fed and formula-fed infants by real-time PCR. FEMS Microbiol Lett 243, 141–147 (2005).

38. Bouillaut, L., Self, W. T. & Sonenshein, A. L. Proline-dependent regulation of Clostridium difficile Stickland metabolism. J Bacteriol 195, 844–854 (2013).

39. Ianiro, G. et al. Variability of strain engraftment and predictability of microbiome composition after fecal microbiota transplantation across different diseases. Nat Med 28, 1913–1923 (2022).

40. Dawkins, J. J. et al. Gut metabolites predict Clostridioides difficile recurrence. Microbiome 10, 87 (2022).

41. Pakpour, S. et al. Identifying predictive features of Clostridium difficile infection recurrence before, during, and after primary antibiotic treatment. Microbiome 5, 148 (2017).

42. Callahan, B. J. et al. DADA2: High-resolution sample inference from Illumina amplicon data. Nat Methods 13, 581–583 (2016).

43. Li, D., Liu, C.-M., Luo, R., Sadakane, K. & Lam, T.-W. MEGAHIT: an ultra-fast single-node solution for large and complex metagenomics assembly via succinct de Bruijn graph. Bioinformatics 31, 1674–1676 (2015).

44. Hyatt, D. et al. Prodigal: prokaryotic gene recognition and translation initiation site identification. BMC Bioinformatics 11, 119 (2010).

45. Buchfink, B., Xie, C. & Huson, D. H. Fast and sensitive protein alignment using DIAMOND. Nat Methods 12, 59–60 (2015).

46. Kanehisa, M. & Goto, S. KEGG: kyoto encyclopedia of genes and genomes. Nucleic Acids Res 28, 27– 30 (2000).

47. Langmead, B., Wilks, C., Antonescu, V. & Charles, R. Scaling read aligners to hundreds of threads on general-purpose processors. Bioinformatics 35, 421–432 (2019).

48. Kang, D. D. et al. MetaBAT 2: an adaptive binning algorithm for robust and efficient genome reconstruction from metagenome assemblies. PeerJ 7, e7359 (2019).

49. Parks, D. H., Imelfort, M., Skennerton, C. T., Hugenholtz, P. & Tyson, G. W. CheckM: assessing the quality of microbial genomes recovered from isolates, single cells, and metagenomes. Genome Res 25, 1043–1055 (2015).

50. Moutinho, T. J., Neubert, B. C., Jenior, M. L. & Papin, J. A. Quantifying cumulative phenotypic and genomic evidence for procedural generation of metabolic network reconstructions. PLoS Comput Biol 18, e1009341 (2022).

51. Orth, J. D., Thiele, I. & Palsson, B. Ø. What is flux balance analysis? Nat Biotechnol 28, 245–248 (2010).

52. Fritzemeier, C. J., Hartleb, D., Szappanos, B., Papp, B. & Lercher, M. J. Erroneous energy-generating cycles in published genome scale metabolic networks: Identification and removal. PLoS Comput Biol 13, e1005494 (2017).

53. Dunphy, L. J., Grimes, K. L., Wase, N., Kolling, G. L. & Papin, J. A. Untargeted Metabolomics Reveals Species-Specific Metabolite Production and Shared Nutrient Consumption by Pseudomonas aeruginosa and Staphylococcus aureus. mSystems 6, e0048021 (2021).

54. Polage, C. R. et al. Overdiagnosis of Clostridium difficile Infection in the Molecular Test Era. JAMA Intern Med 175, 1792–1801 (2015).

55. Dixon, P. VEGAN, a package of R functions for community ecology. Journal of Vegetation Science 14, 927–930 (2003).

56. Genuer, R. & Poggi, J.-M. Random Forests with R. (Springer International Publishing, 2020). doi:10.1007/978-3-030-56485-8.

57. Giorgino, T. Computing and Visualizing Dynamic Time Warping Alignments in *R* : The dtw Package. J. Stat. Soft. 31, (2009).

58. Strobl, C., Boulesteix, A.-L., Kneib, T., Augustin, T. & Zeileis, A. Conditional variable importance for random forests. BMC Bioinformatics 9, 307 (2008).

