## Supplementary figures and images for "Systems-ecology designed bacterial consortium protects from severe *Clostridioides difficile* infection"

### Figure_S1

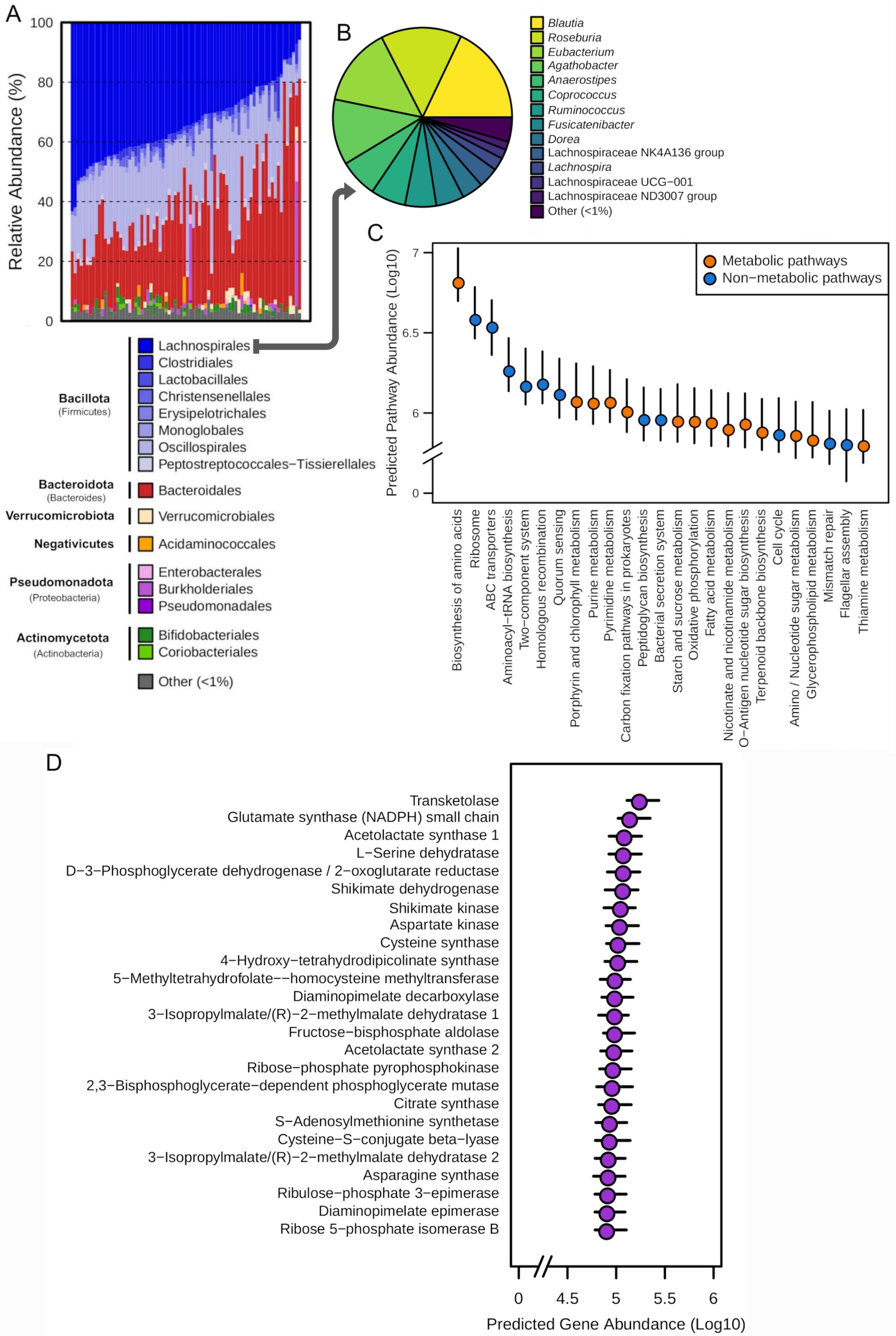

### Figure_S2

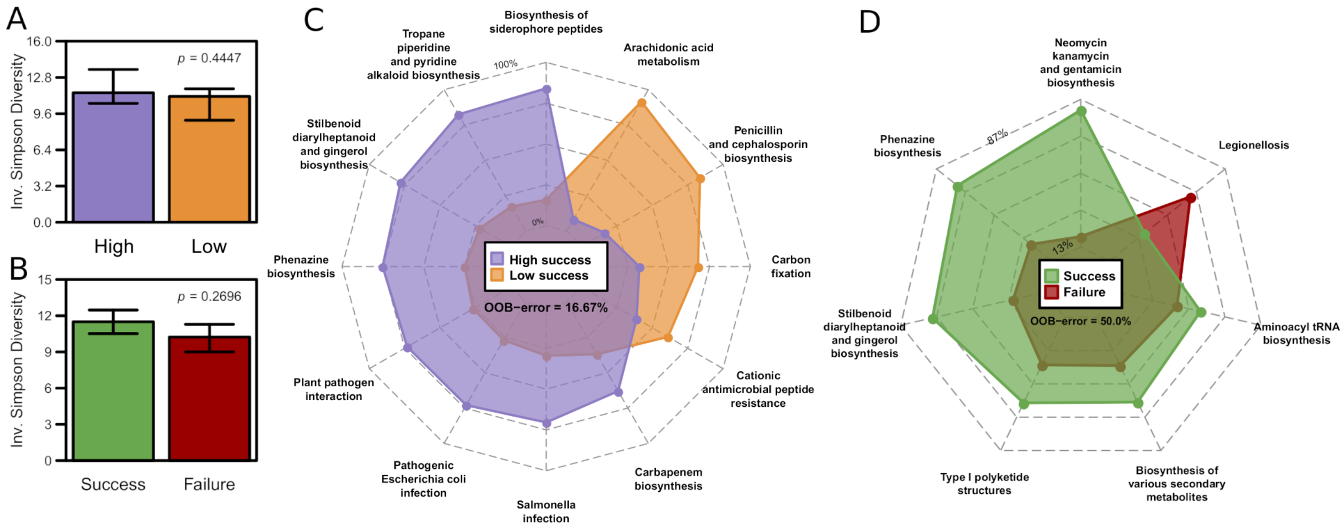

### Figure_S3

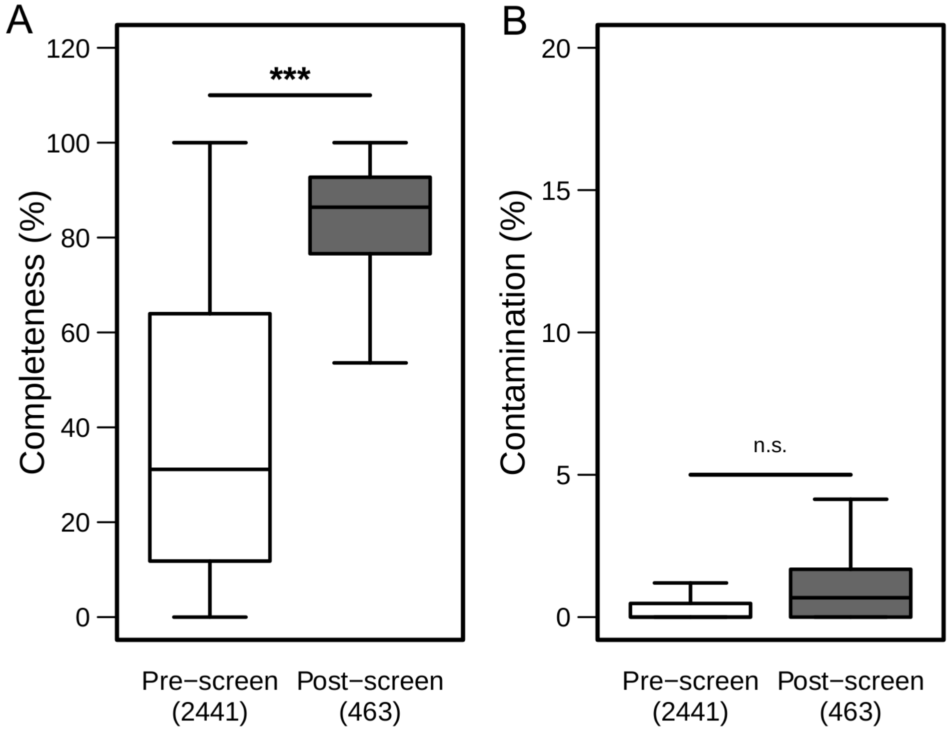

### Figure_S4

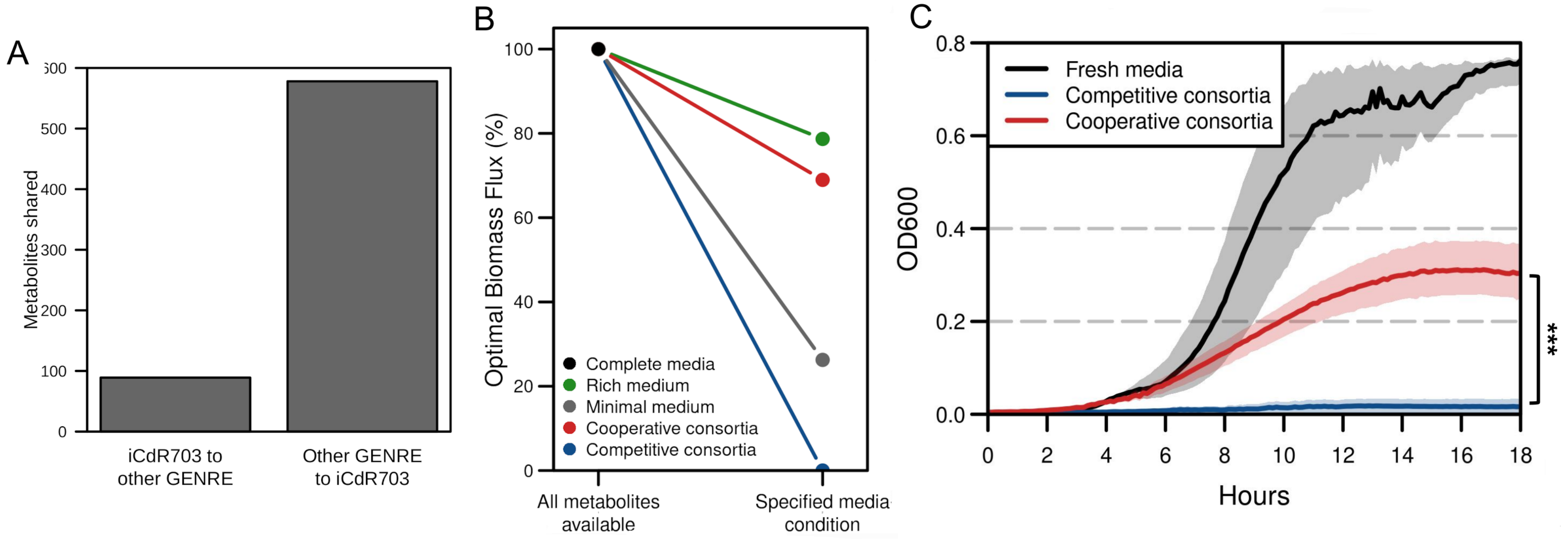

### Figure_S5

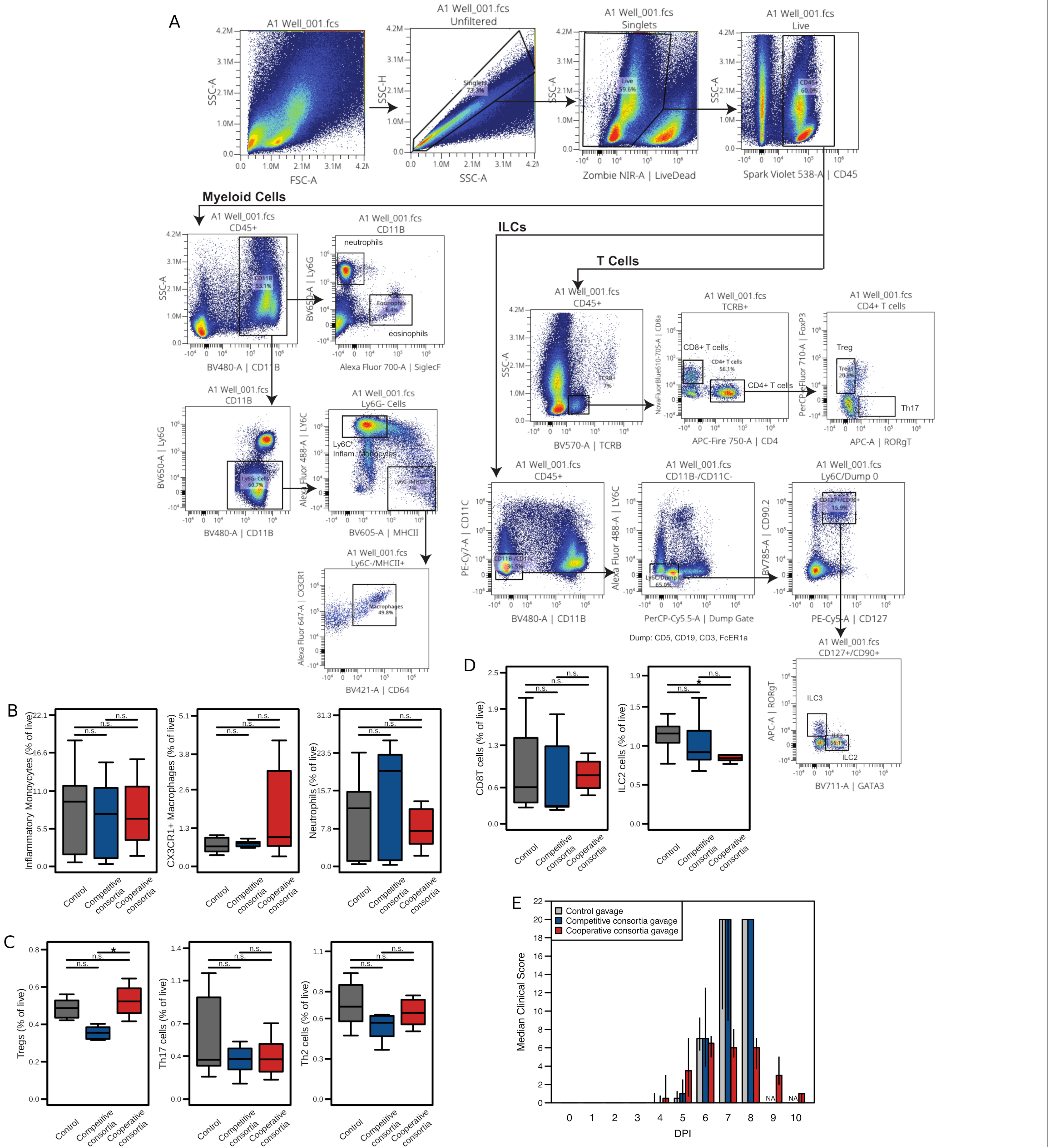

### Figure_S6

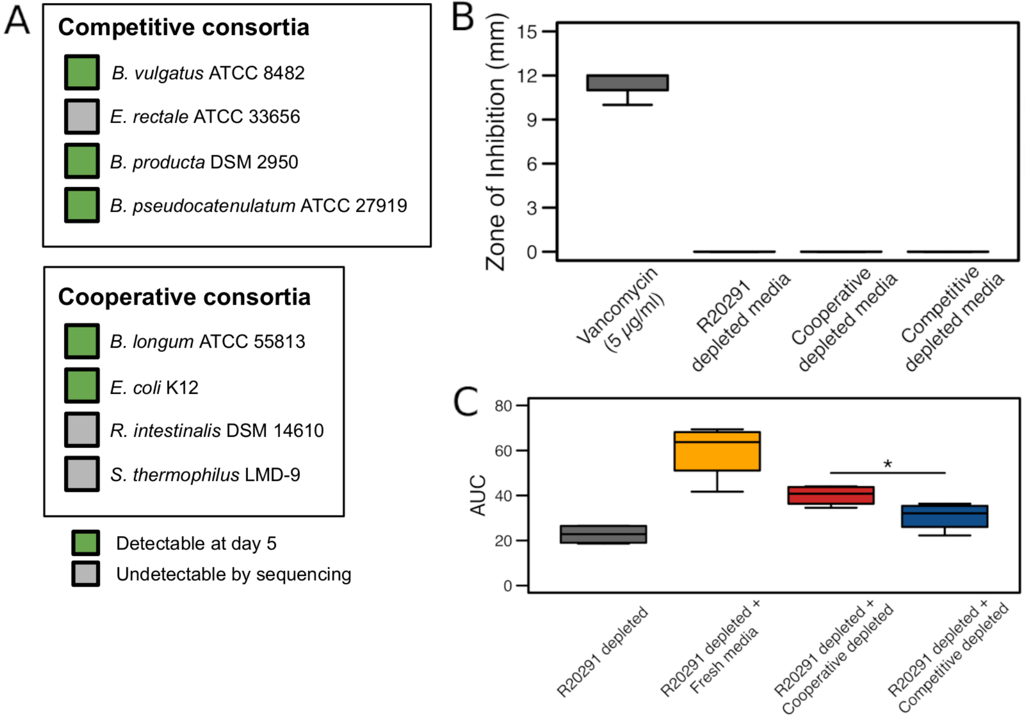
